# Single-molecule tracking reveals the functional allocation, *in vivo* interactions and spatial organization of universal transcription factor NusG

**DOI:** 10.1101/2022.11.21.517430

**Authors:** Hafez el Sayyed, Oliver J. Pambos, Mathew Stracy, Max E. Gottesman, Achillefs N. Kapanidis

**Affiliations:** Gene Machines group, Clarendon Laboratory, Department of Physics, University of Oxford, Oxford, UK; Kavli Institute of Nanoscience Discovery, Dorothy Crowfoot Hodgkin Building, University of Oxford, Oxford, UK; Sir William Dunn School of Pathology, University of Oxford, South Parks Rd, Oxford, UK; Department of Microbiology & Immunology, Columbia University Medical Center, New York, USA

## Abstract

Bacterial gene expression is highly regulated to allow cells to grow and adapt. Much regulation occurs during transcription elongation, where RNA polymerase (RNAP) extends nascent RNA transcripts aided by global and universally-conserved elongation factor NusG. NusG modulates transcription by inhibiting pausing and backtracking; promoting anti-termination on ribosomal RNA (*rrn*) operons; coupling transcription with translation on mRNA genes; and stimulating Rho-dependent termination on toxic genes. Despite extensive work on NusG, its functional allocation and spatial distribution *in vivo* is unknown. Here, we addressed these long-standing questions using single-molecule tracking and super-resolution imaging of NusG in live *E. coli* cells. We found that, under conditions of moderate growth, NusG is mainly present as a population that associates indirectly with the chromosome via RNAP in transcription elongation complexes, and a slowly diffusing population we identified as a NusG complex with the 30S ribosomal subunit; this complex offers a “30S-guided” path for NusG to enter transcription elongation. Only ~10% of total NusG was fast-diffusing, with the mobility of this population suggesting that free NusG interacts non-specifically with DNA for >50% of the time. Using antibiotics and deletion mutants, we showed that most chromosome-associated NusG is involved in *rrn* anti-termination and in transcriptiontranslation coupling. NusG involvement in *rrn* anti-termination was mediated via its participation in phase-separated transcriptional condensates. Our work illuminates the diverse activities of a central regulator while offering a guide on how to dissect the roles of multi-functional machines using in vivo imaging.

## INTRODUCTION

Transcription, a biochemical process central to all living organisms, is orchestrated by the multifunctional protein machine RNA polymerase (RNAP). After the initial recognition of a promoter region, RNAP initiates RNA synthesis and escapes from the promoter after transcribing up to ~15 nt (Mooney et al., 2005; Saecker et al., 2011). RNAP then enters the phase of transcription elongation, which continues processively until termination signals are encountered, at which point the nascent RNA and RNAP dissociate from the DNA (Ray-Soni et al., 2016). During transcription elongation, RNAP encounters DNA sequences and other signals that can slow down or stop transcription via processes such as pausing, backtracking and premature transcription termination. To counteract these impediments, maintain a steady RNA production, and enable additional mechanisms for transcriptional fidelity and regulation, cells employ a large variety of elongation factors that interact in different ways with RNAP.

A key elongation factor, and the only one conserved throughout the tree of life, is bacterial protein NusG, with its archaeal and mammalian homolog being Spt4/5. In *E. coli*, NusG is a 21-kDa protein which consists of two domains connected by a flexible linker; the NusG-NTD domain interacts with RNAP, whereas the NusG-CTD domain has many interacting and mutually exclusive partners (Mooney et al., 2009a). Functionally, NusG appears to play many roles in elongation (Fig. 1A). NusG inhibits transcriptional pausing by enhancing transcription elongation processivity (Kang et al., 2018). NusG also forms an anti-termination complex on ribosomal RNA (rRNA), which maintains steady rRNA transcription with the help of other proteins, including NusE, NusB, ShuB and NusA (Huang et al., 2019, 2020; Krupp et al., 2019; Zellars and Squires, 1999). Further, NusG acts as a “bridge” to couple transcription with translation by linking RNAP to ribosomes via an interaction of NusE/S10 with the NusG-CTD (Bailey et al., 2021; Saxena et al., 2018; Washburn et al., 2020; Webster et al., 2020). Finally, NusG stimulates termination factor Rho, an RNA helicase that terminates the synthesis of toxic gene products associated with homologous gene transfer and dormant phage genes embedded in the bacterial genome (Cardinale et al., 2008; Peters et al., 2009).

**Figure 1.**
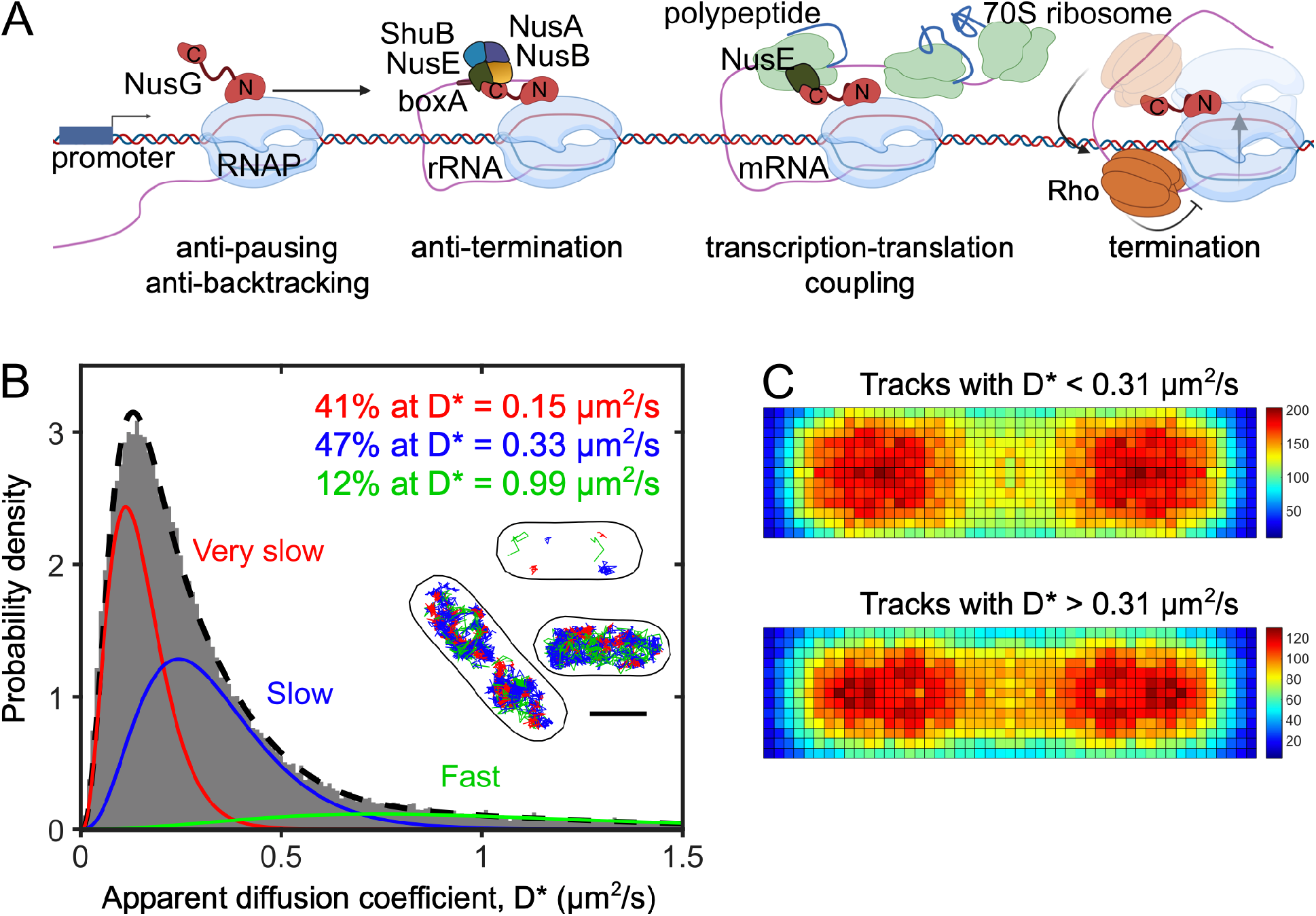
NusG main functions and diffusion landscape in living *E. coli* cells. **A.** Schematic representation of NusG functions and its main interactions during transcription elongation (see text for details). **B.** Distribution of the apparent diffusion coefficients (D*) for 75,264 NusG molecules in live cells grown in M9GluVA. The distribution is best fit by three populations with different mobilities: very-slow NusG (VS-NusG; in red) fixed at D*_vslow_ = 0.15 μm^2^/s; slowly-diffusing NusG (S-NusG, in blue) fitted with a D*_slow_ = 0.33 μm2/s; and fast-diffusing NusG (F-NusG, in green) fitted with D*_fast_ = 0.99 μm2/s. Inset: representative examples of NusG tracks coloured as red, blue, and green for VS-, S-, and F-NusG, respectively. For clarity, only 6 tracks are shown in the top cell. Scale bar, 1 μm. **C**. Spatial distribution heatmaps of NusG tracks with a categorization threshold of D* = 0.31 for 245 cells with lengths of ~2-3 μm, having ~2 nucleoids. Top: a heatmap for molecules with D* < 0.31 μm2/s, representing mainly the VS-NusG species. Bottom: a heatmap for molecules with D* > 0.31 μm2/s, representing mainly the S-NusG and F-NusG species.

Despite extensive study of the roles of NusG in transcription elongation, it is currently unknown how NusG distributes between all these activities in the cell, how the NusG distribution is affected by cell physiology, and whether (or when) NusG constitutes a limiting resource for the cell. This knowledge gap is further exacerbated by the fact that much of our understanding of NusG mechanisms is derived from *in vitro* work, therefore lacking the cellular context, the complexity of the chromosomal structure and organisation, and the presence of competing processes. For example, there is controversy about the exact biological and mechanistic significance of recent cryo-EM structures of large transcription-translation coupled complexes, some of which contain NusG and some do not, and in a way that depends on the length of nascent mRNA emerging from RNAP (Kohler et al., 2017; Webster et al., 2020).

Here, we advance our understanding of NusG functions by examining how NusG distributes between its various activities *in vivo*. To achieve this, we performed single-molecule tracking and super-resolution imaging of NusG fusions with photoactivatable protein PAmCherry, an approach we previously used to study proteins involved in transcription, DNA repair, and chromosome organization (Garza de Leon et al., 2017; Stracy et al., 2015, 2021; Uphoff et al., 2013; Zawadzki et al., 2015). Under conditions supporting moderate growth rates, we found that most of NusG is present in a chromosome-associated population (via an elongating RNAP) and a slowly-diffusing population corresponding to a NusG complex with free 30S ribosomal subunit; notably, there is little free NusG in the cell. Using mutants of NusB and of NusE, we show that most chromosome-associated NusG is involved in anti-termination on rRNA genes, and in transcription-translation coupling. Finally, using a chemical treatment that removes clusters arising from liquid-liquid phase separation (LLPS), we show that NusG clusters are associated with LLPS-driven condensates involved in rRNA anti-termination.

## RESULTS

### Construction and characterization of a functional NusG-PAmCherry fusion

To study the spatial distribution and mobility of NusG in bacterial cells, we first constructed a functional NusG-PAmCherry fusion in *E. coli* strain MG1655, where *nusG* is part of the essential *secE-nusG* operon. We first attempted to insert PAmCherry at the C-terminus of NusG using λ red recombination (Datsenko and Wanner, 2000), but no viable fusions were obtained. We then inserted a PAmCherry, along with a flexible linker, at the NusG N-terminus; since this strategy required inserting the PAmCherry gene between *secE* and the *nusG* coding sequence (where having an antibiotic marker for successful integration was not possible), we used gene gorging (Herring et al., 2003; Lee et al., 2009; Mooney et al., 2009b). The resulting strain carrying NusG-PAmcherry exhibited normal growth compared to the WT (Fig. S1A); further, the fusion was expressed as an intact protein (Fig. S1B).

### Single-molecule tracking reveals a wide range of NusG mobilities *in vivo*

To study NusG mobility and spatial distribution in live cells, we used single-molecule tracking combined with photoactivation localization microscopy (tracking-PALM; (Manley et al., 2008)). Since NusG is a transcription elongation factor involved in many functions occurring on the bacterial chromosome, we expected NusG to distribute in at least two diffusive subpopulations; a very-slowly-moving species representing NusG molecules that bind stably with RNAP during transcription elongation on the chromosome (and therefore adopt the very low mobility of the chromosomal loci), and a fast-moving species representing free NusG molecules (~50 KDa for the entire fusion construct). In our previous work on RNAP, we showed that tracking PALM is able to capture this entire range of intracellular mobilities (Stracy et al., 2015).

We first studied NusG mobility in M9 minimal media supplemented with glucose, MEM vitamins and MEM amino acids (hereafter, “M9GluVA”). Using photoactivation and imaging using 10.64-ms exposures, we collected 75,264 single-molecule tracks, calculated their apparent diffusion coefficient (*D*\*), and summarized the mobility of all tracks in a D* distribution (gray histogram, Fig. 1B). The D* distribution showed that the large majority of molecules appear to have fairly low mobility (D*<0.5 μm^2^/s). As with other DNA-binding proteins (Stracy et al., 2015, 2016; Uphoff et al., 2013), the shape of the distribution is complex, and could not be fit well by a single diffusing species (Fig. S2A). A two-species free fit described the distribution better (Fig. S2B), splitting approximately evenly between a very-slow species (D*_vslow_~0.15 μm^2^/s) and a slow species (D*_slow_~0.4 μm^2^/s).

To explore the possibility that our fitting above fails to capture a minor fast-diffusing species (with D*~1 μm^2^/s), we also performed a three-species fit, with the D* of the very-slow species fixed to the value obtained by the two-population fit. The three-species fit fitted our distribution extremely well (Fig. 1B), and showed, in addition to the two main species, the presence of a minor fastdiffusing species (D*_fast_~1.0 μm2/s) that accounts for ~12% of all NusG. Both two- and three-species fits showed that there is little fast-diffusing NusG in the cell, a species which we will refer to as “free NusG” or F-NusG.

We then tentatively assigned the mobility species to different activities or complexes of NusG. Since the very-slow species (~41% of all tracks; hereafter, the “VS-NusG” species) has a very low mobility, similarly to proteins that bind stably to the chromosome (Stracy et al., 2015), we initially assigned this species to NusG molecules indirectly (and stably) associated with the chromosome via interactions with RNAP and, possibly, via other machinery that interacts with the chromosome during active elongation, anti-termination, or Rho-dependent termination.

Intriguingly, however, the largest fraction (~47%) of the tracks is due to the slow species (D* ~0.3 μm^2^/s; hereafter, the “S-NusG” species). This mobility is much slower than that expected for free NusG, indicating that S-NusG comprises NusG molecules with slowed-down motions due to interactions with much larger structures. The structures are likely to be one or more of the three major interacting partners of NusG in elongation: the RNAP core, the Rho-termination factor, and the ribosome or ribosomal subunits.

We also found that the distribution between the three main diffusive species in M9GluVA was similar to that in a medium supporting significantly slower growth (M9 + 0.2% glucose medium, hereafter “M9Glu”, with a generation time of ~55 min; Fig. S2D), as well as to a medium supporting significantly faster growth (rich-defined medium, hereafter “RDM”, with a generation time of ~33 min).

To map the sub-cellular distributions of the NusG species and test our initial assignments, we divided the NusG tracks into a fraction comprised mainly by VS-NusG, and a fraction comprised mainly by S-NusG (by selecting tracks with D*<0.31, and D*>0.31, respectively), and plotted heatmaps for the tracks of these two fractions (Fig. 2C and Fig. S2E, for cell-size ranges featuring two nucleoids vs. one nucleoid, respectively). The VS-NusG distribution resembled the distribution of chromosome-engaged RNAPs, which tend to localize at the periphery of the nucleoid (Stracy et al., 2015), and is consistent with the assignment VS-NusG to NusG molecules engaged in transcription elongation.

**Figure 2.**
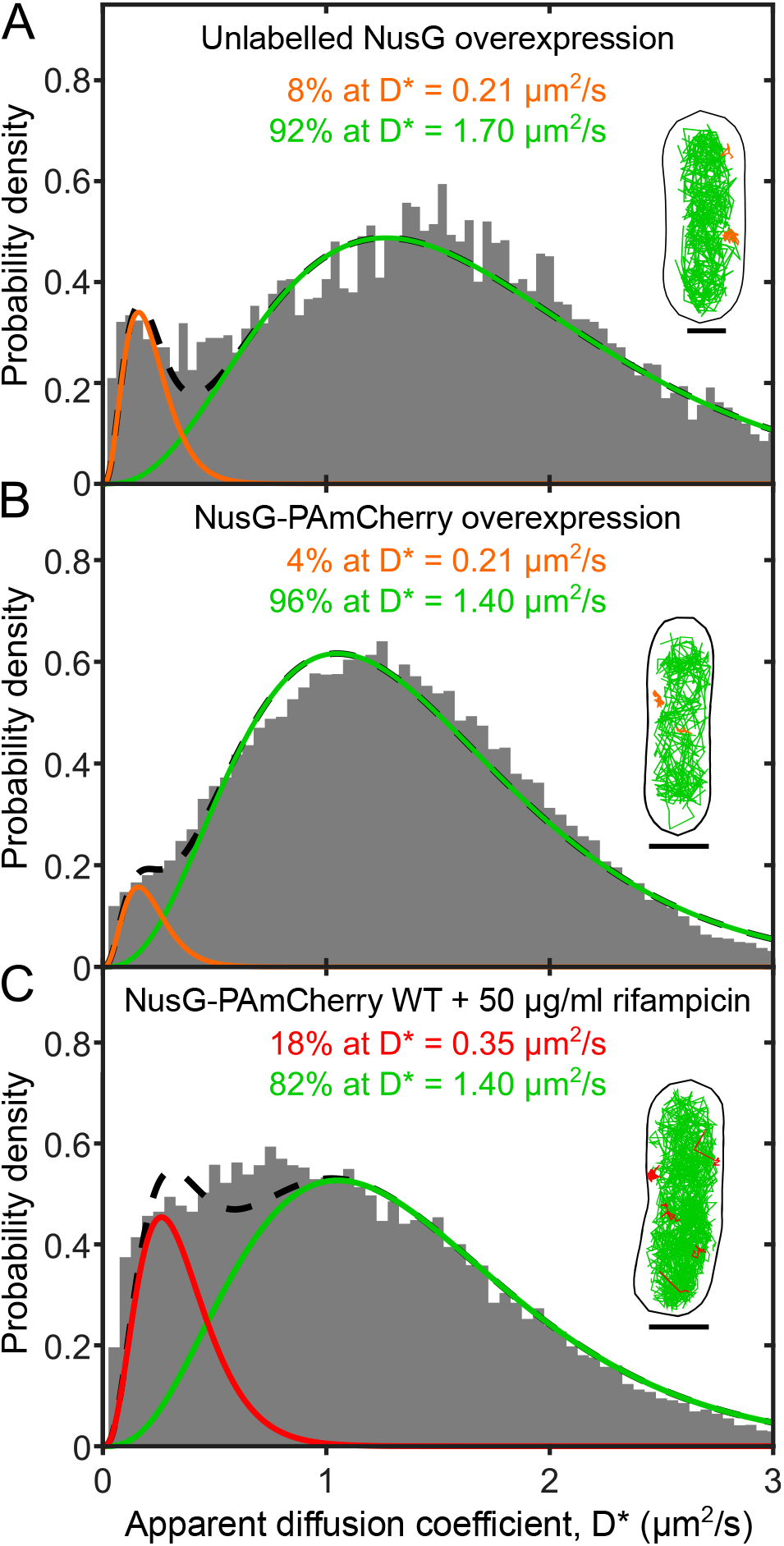
Measuring the intracellular mobility of fast-diffusing NusG in M9GluVA. **A.** D* distribution for 9,751 NusG molecules in live cells after overexpressing unlabeled NusG for 30 min post induction with 1 mM IPTG. The distribution is best fit by two populations with D* of 0.21 μm2/s and 1.7 μm2/s. Inset, representative examples of tracks corresponding to the two populations; the categorization threshold was 0.25 μm2/s. Scale bar, 1 μm. **B.** D* distribution for 62,867 NusG molecules in live cells after overexpressing NusG-PAmCherry for 30 min post induction with 1 mM IPTG. The fit and representative tracks were prepared as in panel A. **C.** D* distribution for 28,198 NusG molecules in live cells after inhibiting transcription initiation using 50 μg/ml rifampicin (Rif) for 30 min. The fit and representative tracks were prepared as in panel A, with the exception of the D* of the F-NusG being fixed at D*_fast_ =1.4 μm2/s, and the categorization threshold for track colouring being 0.35 μm2/s.

In contrast, the S-NusG fraction is found throughout the cytoplasm and not exclusively in the nucleoid. This localization pattern is different from the localization of diffusing RNAP, which we had previously shown to localize almost exclusively to the nucleoid region due to non-specific interactions with the chromosome (Bakshi et al., 2012; Stracy et al., 2015). These results strongly suggest that the S-NusG is formed due to a NusG interaction with a large structure other than RNAP. Further, our results are consistent with an interaction of NusG with the 30S free ribosomal subunit, since the latter localizes throughout the cell, and is not excluded from the nucleoid (Sanamrad et al., 2014).

### Characterising the fast-diffusing NusG species

Our initial analysis showed that most NusG molecules interact with larger partners, leaving little free NusG (F-NusG) in the cell. To verify this observation, and determine more accurately the mobility of F-NusG, we overexpressed unlabeled NusG from an IPTG-inducible promoter ((Mooney et al., 2010) and *Methods*) to out-compete NusG-PAmCherry from its interactions with its partners, and release the fusion molecules in the cytoplasm.

Indeed, upon overexpression of unlabelled NusG following 30 min induction, the NusG mobility distribution (Fig. 2A) showed near complete disappearance of VS-NusG and S-NusG (<10% for the sum of the two species). Further, a fast-diffusing species (D*_fast_~1.7 μm^2^/s) became predominant, accounting for >90% of all NusG. These results showed that overexpressed NusG fully replaces the tagged version in its interactions in the cell, and provides a better estimate (due to much better sampling than that in Fig. 1B) of the mobility of F-NusG.

To further validate the mobility of F-NusG without relying on releasing NusG from its complexes, we also overexpressed a NusG-PAmCherry fusion from a low-copy-number plasmid with an IPTG-inducible *plac* promoter in wild-type MG1655. After 30 min of induction, NusG appears almost exclusively (96%) as F-NusG species (Fig. 2B). This experiment also provided an additional estimate for the mobility of F-NusG (D*~1.4 μm^2^/s).

To further support the analysis of above, and confirm whether VS-NusG indeed comprises NusG molecules bound to chromosome-bound RNAP during transcription elongation, we treated cells with the antibiotic rifampicin (Rif) to block initial transcription and subsequent elongation (Campbell et al., 2001; Herring et al., 2005), and determined NusG mobility (Fig. 2C). As with the NusG-overexpression experiments, VS-NusG essentially disappears as a result of Rif treatment, showing that chromosomal association of NusG requires, as expected, entry of RNAP into transcription elongation (Mooney et al., 2009b; Stracy et al., 2015). The Rif treatment also substantially increased the abundance of F-NusG (from 12% to 85% of the entire NusG pool), as was seen after NusG overexpression (Fig. 2A-B). Notably, the S-NusG species was reduced but not abolished by Rif treatment (still accounting for ~18% of all NusG), showing that its presence is not dependent on active transcription.

### Most chromosome-associated NusG is engaged in rRNA antitermination

We then examined what fraction of NusG engages in transcription anti-termination. Processive anti-termination is mediated by a complex of proteins interacting with RNAP to counteract premature termination (Huang et al., 2020; Torres et al., 2004), and ensure efficient *rrn* operon transcription. This is accomplished by the formation of an *rrn* anti-termination complex (ATC), which includes NusA, NusB, NusE, NusG, and ShuB. ATC formation is initiated by the NusB:NusE heterodimer binding to a *boxA* site at the leader rRNA sequence (Fig. 3A; see also (Burmann et al., 2009; Huang et al., 2019, 2020; Krupp et al., 2019)). NusB is found at copy numbers that are 50-80% of those of core RNAP in the cell (Swindle et al., 1988). If NusB is deleted, the assembly of ATC on *rrn* operons is blocked, reducing the total RNA content in the cell and increasing Rho-dependent termination (Miyashita et al., 1982). We thus reasoned that, by eliminating NusB, we can prevent anti-termination, and use the ensuing change in NusG mobility to estimate the fraction of NusG is involved in antitermination.

**Figure 3:**
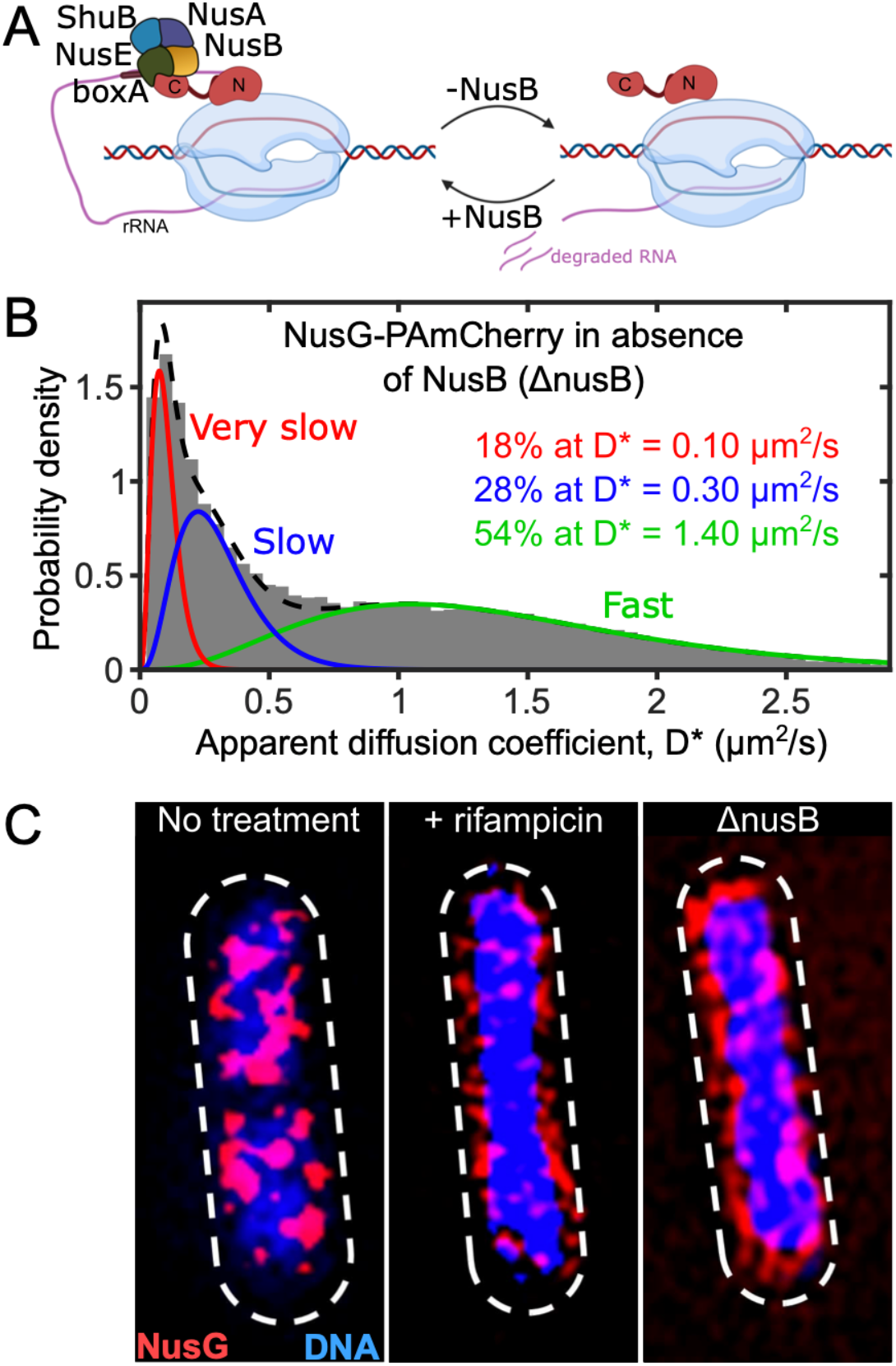
Blocking anti-termination complex formation increases NusG diffusion and alters its spatial distribution. **A.** Schematic depicting the effects of *nusB* deletion (*ΔnusB*) on *rrn* anti-termination. NusB deletion blocks anti-termination complex formation, depletes NusG from *rrn* operons, and leads to premature transcription termination and increased RNA degradation. **B.** D* distribution for 41,524 NusG molecules in live cells deficient in anti-termination complex formation due to *nusB* deletion; the growth medium is M9GluVA. The distribution is best fit by three populations with D*_vslow_ =0.1 μm2/s (in red), D*_slow_ = 0.3 μm2/s (in blue), and D*_fast_ = 1.4 μm2/s (in green). Scale bar, 1 μm. **C.** 3D-SIM imaging of NusG-sfGFP (in red) and DNA stained with DAPI (in blue) for untreated live cells, Rif-treated cells, and *ΔnusB* cells, showing the relative NusG spatial distribution in relation to the nucleoid, and highlighting how both Rif treatment and NusB deletion affect the NusG distribution and nucleoid decompaction. Dashed line: cell outlines.

To study NusG diffusion in the absence of NusB, we studied the NusG mobility distribution in a Δ*nusB* strain (Fig. 3B). NusB deletion resulted in a large increase in F-NusG (which reached 55% of all NusG), in large part due to a ~60% decrease in VS-NusG (from 41% to 17%). This fraction (60%) serves as a lower bound of the fraction of chromosome-associated NusG that is involved in *rrn* anti-termination, since some VS-NusG freed from anti-termination may be directed to other (non-*rrn*) chromosome-associated species. Notably, the S-NusG species remains significant, albeit reduced by ~40%. This reduction is consistent with our assignment of S-NusG to a NusG-30S complex, since loss of rRNA transcription will also reduce the levels of the 30S ribosomal subunit, and, in turn, the levels of the proposed NusG-30S complex; however, these results do not exclude the presence of a putative NusG-Rho complex in the S-NusG species.

We also examined the spatial distribution of the NusG fraction involved in anti-termination using 3D structured illumination microscopy (3D-SIM; see *Methods*). Using a NusG-sfGFP strain grown in M9GluVA, we observed that NusG localised in large clusters (10-20 clusters per cell) that decorate an irregular nucleoid (blue density; Fig. 3C, left). This distribution was similar to that of RNAP in rich media, as we had previously observed using SIM (Stracy et al., 2015). In contrast, Rif treatment removed all large NusG clusters, leaving mainly small regions of nucleoid-peripheral NusG signal over a decondensed nucleoid (Fig. 3C, middle).

NusB deletion also resulted in loss of most large NusG clusters, decondenses the nucleoid, and leads NusG to a more regular nucleoid-peripheral localisation (Fig. 3C, right). The change in the NusG spatial distribution is consistent with the expectation that absence of *rrn* anti-termination will lead to a nucleoid-wide loss of the high levels of transcription of *rrn* operons. The remaining clusters are likely to represent NusG attached to clusters of mRNA-transcribing RNAPs, which should be less affected by the loss of *rrn* anti-termination.

### Loss of NusG contacts with the 30S ribosome release NusG from elongation complexes and the S-species

We then examined the involvement of NusG in transcription-translation coupling, a process extensively studied *in vitro* via biochemical assays and structural approaches (Burmann et al., 2010; Kohler et al., 2017; Washburn et al., 2020; Webster et al., 2020).

To dissect the role of NusG-ribosome interactions mediated via the NusE/S10 subunit on the 30S ribosome (Fig. 4A), we inserted a degron tag in the NusE C-terminus (see *Methods*). NusE interacts with NusG both during transcription-translation coupling, and during the assembly of the *rrn* anti-termination complex. As a result, once NusE is degraded, we expected a large increase in F-NusG; further, if indeed NusE interacts with NusG in the context of the putative NusG-30S complex, we expected S-NusG species to decrease substantially upon NusE degradation.

**Figure 4.**
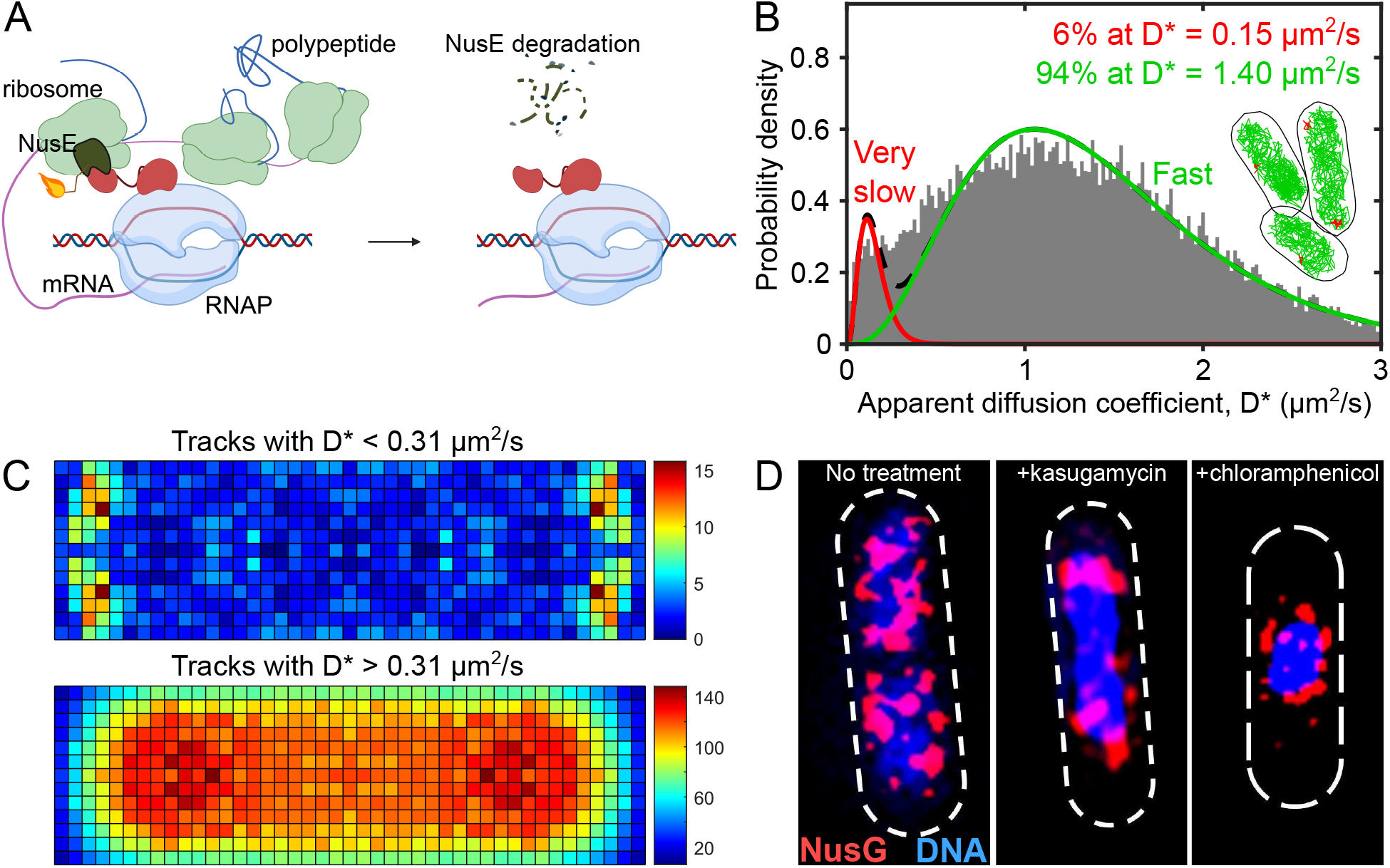
Breaking ribosome-NusG interactions abolishes transcription-translation coupling and eliminates the S-NusG species. **A.** Schematic of interactions between a lead ribosome and NusG in the context of transcription elongation; the interactions are between the ribosomal subunit NusE/S10 and the C-terminal domain of NusG. The fuse depicts the degron tag that leads to NusE degradation (depicted as dashed line in the schematic on the left) upon IPTG addition. **B.** D* distribution for 20,593 NusG molecules in live cells after induction of NusE degradation (1 hr in liquid media, as well as during imaging); the growth medium is M9GluVA. The distribution is best fit by two populations with D*_vslow_ = 0.15 μm2/s (in red), and D*_fast_ = 1.4 μm2/s (in green). Inset, representative examples of tracks corresponding to the two populations (categorization threshold of D* = 0.31). Scale bar, 1 μm. **C.** Spatial distribution heatmaps of NusG tracks with a categorization threshold of D* = 0.31 for 311 cells with lengths of ~2-3 μm, having ~2 nucleoids. Top: a heatmap for molecules with D* < 0.31 μm2/s, representing the VS-NusG species. Bottom: a heatmap for molecules with D* > 0.31 μm2/s, representing the F-NusG species. **D.** 3D-SIM imaging of NusG-sfGFP (in red) and DNA stained with DAPI (in blue) for untreated live cells, cells treated with 500 μg/ml translation initiation inhibitor kasugamycin, and 100 μg/ml translation elongation inhibitor chloramphenicol. Dashed line: cell outlines.

Indeed, upon induction of NusE degradation, the sum of VS-NusG and S-NusG populations was reduced from ~90% to less than 10%, with the large majority of NusG converted to F-NusG (D*~1.4 μm^2^/s; Fig. 4B). These results strongly support our proposal that the S-NusG species corresponds mainly to a NusG-30S complex, and not to a NusG-Rho complex or to an NusG-RNAP, both of which should have been either unaffected or increased upon NusE degradation.

To explore further the effects of NusE degradation on NusG functions, we also examined the NusG spatial distribution after NusE degradation (Fig. 4C), and established that the F-NusG population distributes across the cytoplasm (Fig. 4C, bottom). In sharp contrast, the remaining low-mobility species are completely excluded from the nucleoid periphery (Fig. 4C, top); these species are likely to represent genes actively being prematurely aborted by Rho, as well as any remaining NusG-RNAP complexes.

We then used 3D-SIM to determine the NusG sub-cellular distribution after destabilizing the NusG interactions with RNAP and ribosomes by blocking different steps in translation, while not affecting *rrn* antitermination directly. We reasoned that if NusG couples RNAP with the leading ribosome, blocking translation would remove NusG-ribosome interactions, and thus decrease the overall affinity of NusG for the elongation complex on mRNA genes. Previous work on the spatial distribution of ribosomes relative to the nucleoid had shown that treatment with chloramphenicol (Cam), a translation-elongation inhibitor, leads to filling of most cytoplasmic space with ribosomes, with the nucleoid becoming condensed into a spherical object appearing at mid-cell (Bakshi et al., 2014). Conversely, kasugamycin (Kas), a translation-initiation inhibitor, inhibits 70S ribosome assembly while allowing the nucleoid to maintain most of its length (Bakshi et al., 2014).

After Kas treatment, we observed that medium-size NusG clusters, previously distributed peripherally and along the nucleoid, were lost (cf. Fig. 4D, left with Fig. 4D, middle). In contrast, the large NusG clusters located at the nucleoid edges closest to the cell poles remained unaffected, and are likely to reflect NusG bound to *rrn* operons. On the other hand, Cam treatment led to a spherical nucleoid surrounded by NusG clusters (Fig. 4D, right); this NusG localisation resembled closely the RNAP distribution seen in cells treated with Cam (Bakshi et al., 2012; Stracy et al., 2015).

### A large fraction of NusG is confined in condensates formed via liquid-liquid phase separation

Our NusG diffusion analysis showed that deletion of *nusB* (Fig. 3B) and consequent failure to assemble *rrn* anti-termination complexes resulted in a large decrease (~60%) in the abundance of VS-NusG. This decrease was also accompanied by a significant and unexpected decrease in the mobility of the remaining VS-NusG from a D*~0.15 μm^2^/s to ~0.10 μm^2^/s (Fig. 3B).

To understand the origin of this decreased mobility, we considered the findings of Ladouceur and coworkers (Ladouceur et al., 2020), where *nusB* deletion led to loss of large RNAP clusters *in vivo*. The same work showed that anti-termination factor NusA can form phase-separated liquid droplets *in vitro*, can drive foci formation *in vivo*, and may drive LLPS of RNAPs and components of the *rrn* anti-termination complex *in vivo*. To test for the presence of condensates, Ladouceur et al. treated cells with 1,6-hexanediol (HEX), an aliphatic alcohol that destabilizes liquid condensates but not protein aggregates (Kroschwald et al., 2017). HEX exposure induced loss of RNAP and NusA clustering, providing strong evidence for RNAP/NusA LLPS *in vivo*.

To test whether NusG clustering is linked to RNAP/NusA condensates, we grew cells expressing NusG-sfGFP in M9GluVA, immobilized them on agarose pads in the absence or presence of 5% HEX, incubated for 5 min, and imaged the cells (Fig. 5A, top panels). Untreated cells showed large NusG clusters in the form of bright spots within the cytoplasm, whereas NusG-sfGFP clusters disappear in the presence of HEX. We repeated this comparison after treatment with Kas in liquid for 30 min, a treatment expected to retain mainly the NusG clusters involved in *rrn* antitermination complexes (Fig. 4D) before immobilization on agarose. As predicted, Kas-treated cells displayed the NusG clusters more clearly (Fig. 5A, bottom left), while subsequent HEX treatment greatly diminished NusG clustering. These results strongly support that there is a substantial presence of NusG in LLPS-driven condensates of the *rrn* anti-termination machinery *in vivo*.

**Figure 5.**
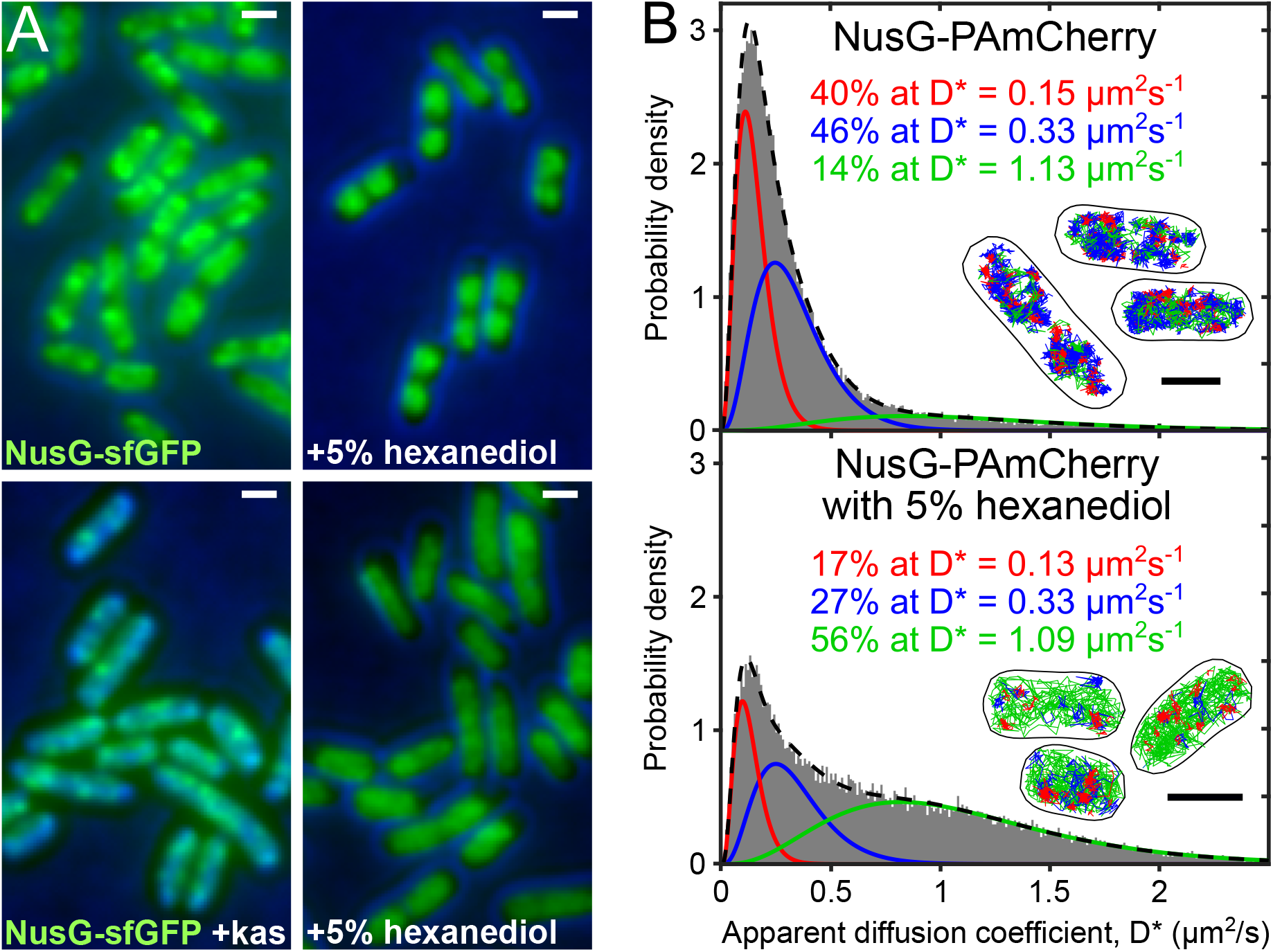
Treatment with 1,6-hexanediol eliminates NusG clusters and releases NusG from biomolecular condensates involved in transcription anti-termination. **A.** NusG-sfGFP images of live cells imaged using 10-ms exposures. Results obtained from cells grown in richM9 media until OD600 ~ 0.2, and split into cells that remain untreated and cells that were treated for 30 min in liquid culture with kasugamycin. Both untreated and Kas-treated cells were then immobilized on agarose pads with or without 5% 1,6-hexanediol (HEX) for 5 min prior to imaging. **B.** D* distribution for live cells containing NusG-PAmCherry in the absence (top) and presence of 5% HEX for 5 min (bottom). The D* for untreated cells is similar to that in Fig. 1B; the D* distribution for HEX-treated cells is best fit by three populations with D*_vslow_ = 0.13 μm2/s (in red), D*_slow_ = 0.33 μm2/s (fixed; in blue), and D*_fast_ = 1.09 μm2/s (in green). Inset, representative examples of tracks corresponding to the three populations; the categorization thresholds were D* < 0.2 μm2/s for the red tracks, 0.2 < D* < 0.5 μm2/s for the blue tracks, and D* > 0.5 μm2/s for the green tracks. Scale bar, 1 μm.

We also examined the effect of HEX treatment on NusG mobility by performing tracking PALM on HEX-treated cells expressing NusG-PAmCherry in M9GluVA (Fig. 5B). Treatment by HEX dramatically changed the NusG diffusion profile. First, the abundance of VS-NusG decreased by ~60%, from 41% in untreated cells to 17% in HEX-treated cells. Second, the abundance of F-NusG species increased dramatically, from 12% in untreated cells to 56% in HEX-treated cells. Third, the abundance of S-NusG also decreased by ~40%.

These striking results suggest that, under moderate growth conditions, ~25% of all NusG is confined in condensates. The results were also consistent with the *ΔnusB* results (Fig. 3B), which showed an identical decrease in the abundance of VS-NusG and S-NusG species, as well as a similar reduction in the mobility of the VS-NusG. Taken together, the results of the HEX treatment and the *nusB* deletion clearly establish that these conditions disrupt the condensates in a similar fashion, and free the condensate-confined pool of NusG.

## DISCUSSION

Here, we combine our powerful single-molecule tracking approach with mutational analysis *in vivo* to map the spatial distribution and dissect the functions of transcription elongation factor NusG using *E. coli* as a model organism. Our analysis complements the extensive *in vitro* analysis of NusG activities and interactions using structural, biochemical, and biophysical approaches, as well as *in vivo* analysis using genetic and cell-biology approaches. Our results offer direct views of the allocation of the NusG pool between its many functions, and provide estimates that can help model transcription and gene expression in bacterial cells. Since NusG is the only transcription factor conserved amongst all kingdoms of life, many of our conclusions are likely to hold true for many organisms other than *E. coli*. Our approach also provides a roadmap on how to analyse activities of multi-functional proteins *in vivo* using imaging.

### Only ~10% of NusG is free in the cell

NusG is a multi-functional global elongation factor involved in many complexes nucleated on RNAP molecules during transcription elongation. It was unknown, however, what fraction of NusG is involved in such chromosome-associated complexes under different growth conditions. Our work shows that for media supporting doubling times of 33-55 min, there is a limited amount of free NusG (corresponding to the F-NusG species) in the bacterial cytoplasm, with only ~10% of NusG appearing to diffuse rapidly. This result strongly suggests that NusG is a limited resource for the cell, and predicts the presence of a dynamic competition between machineries for binding NusG, with the split between activities changing according to the cellular requirements for growth, duplication, and adaptation.

### Free NusG binds to the chromosome non-specifically and transiently for >50% of time

Our work clearly established that free NusG-PAmCherry, a protein of ~48 KDa, has an apparent diffusion coefficient of D*_fast_ ~ 1.4 μm^2^/s (Fig. 2). Given that NusG is not known to bind directly to chromosomal DNA, but instead binds to the chromosome indirectly (via RNAP during transcription elongation), we would expect that the D* value and diffusion behaviour of NusG should be independent of the presence of the chromosome, as we have shown for proteins lacking DNA-binding domains (Stracy et al., 2021). However, the D* value for free NusG matches that for a 4-times larger protein that is unable to bind DNA (Lac^-41^, a ~200 KDa truncated *lac* repressor (LacI) derivative lacking its DNA-binding domain).

Further, under the same consideration that NusG does not bind directly to chromosomal DNA, the diffusion of free NusG in cells with intact nucleoids should resemble the behaviour of HU-PAmCherry, 48 KDa (same size as NusG-PAmCherry) in cells where the chromosomal DNA has been degraded (DNA-free cells; see (Stracy et al., 2021)). Again, strikingly, while the estimated accurate diffusion coefficient of HU-PAmCherry in DNA-free cells is D_acc_ ~12.6 μm^2^/s (Stracy et al., 2021), the D_acc_ for NusG-PAmCherry in cells with intact nucleoids is estimated to be ~3.5 μm^2^/s (based on the similarity of the D* values for NusG-PAmCherry and LacI-PAmCherry, and the conversion of D* to D_acc_ for LacI-PAmCherry using simulations of diffusion (Stracy et al., 2021)).

These results strongly suggest that either NusG interacts in the cell with another diffusing biomolecule such as a high-copy number protein, e.g., NusA, shown to interact *in vitro* with NusG (Strauß et al., 2016), or that NusG binds non-specifically and transiently to chromosomal DNA. The strong nucleoid-like localisation of F-NusG (Fig. 2B-C) supports the second hypothesis. Further, the fact that the F-NusG species persists and shows no change in mobility in rifampicin-treated cells, where RNA is highly depleted, indicates that these non-specific interactions are not mediated primarily by nascent RNA. As a result, we can use the estimated D_acc_ for NusG in the absence and presence of the nucleoid (~12.6 μm^2^/s and ~3.5 μm^2^/s, respectively) to estimate that free NusG binds to the chromosome transiently for ~70% of its diffusion time (see *Methods*). We note that interactions between the DNA and NusG-like proteins, especially with non-template DNA in the context of the transcription bubble, have been previously discussed (Nedialkov et al., 2018). The non-specific interactions of NusG with the chromosome may increase the effective concentration of NusG in the vicinity of elongation complexes, thus facilitating the search of NusG for some of its targets.

### NusG interacts with the 30S ribosomal subunit before translation initiation

Our work establishes that most NusG associates with transcription elongation complexes on the chromosome (forming VS-NusG species), or diffuses as part of larger complexes (i.e., S-NusG species). Notably, NusG binds to the nucleoid stably only in the presence of transcription elongation, as shown by Rif treatment (Fig. 2C); this is further supported by the similarity of the spatial distributions of VS-NusG (Fig. 1C, top) with chromosome-associated RNAP (Stracy et al., 2015).

Our detection of an abundant S-NusG species is intriguing. Our NusE-degradation results (Fig. 4) ruled out free RNAP and Rho as large protein partners of NusG in S-NusG. We also rule out the fully assembled 70S ribosome as the potential partner, since the slow-diffusing species has full access to the nucleoid, in contrast to the 70S ribosome, which does not enter the nucleoid in a diffusing form (Sanamrad et al., 2014).

Instead, the only large complex that can access the nucleoid and interact with NusG is the 30S ribosomal subunit, which is not excluded from the nucleoid and which has a diffusion coefficient similar to that we observe here (D*~0.3-0.4 μm^2^/s for 30S; (Bakshi et al., 2012, 2014; Sanamrad et al., 2014)). Our NusE-degradation results further support the presence of NusG-30S complex, since loss of NusE essentially eliminates S-NusG. Consistent with this interpretation, structural studies suggest that NusG:NusE interaction occurs both in the context of the 70S ribosome, and in the context of free 30S (Burmann, et al., 2010).

The presence of substantial amounts of a NusG-30S complex also suggests an additional major route of NusG entry to the transcription elongation complex – that of association during translation initiation through the interaction of a NusG-loaded 30S with the Shine-Dalgarno sequence on mRNA, formation of 70S ribosomes, ribosome translocation towards an elongating (or stalled) RNAP, and NusG-facilitated transcription-translation coupling on target mRNA genes. This is in line with the ChIP-chip analysis (Mooney et al., 2009b) showing that NusG enters the transcription elongation complex on mRNA a few hundred bp from the transcription start site (Mooney et al., 2009b). This “30S-guided” mode of entry is in addition to a simpler mode where NusG enters transcription elongation by free NusG binding directly to RNAP molecules after they enter elongation, with an *in vitro* measured K_d_ of ~120 nM; (Turtola and Belogurov, 2016)).

### Most NusG is involved in transcription anti-termination on rRNA genes and in transcription-translation coupling

NusG is known to interact with NusB, NusE and other proteins to form an anti-termination complex on *boxA* sequences at the 5’ end of rRNA, and in turn, protect rRNA from premature termination (Peters et al., 2009; Torres et al., 2004). Since NusB:NusE dimerization on *boxA* is the pre-requisite for anti-termination complex assembly, *nusB* deletion results in 95-97% reduction in anti-termination in rRNA genes, the expression of which accounts for the majority of RNA in cells growing at moderate growth rates (Bremer and Dennis, 2008).

Our mobility analysis showed that *nusB* deletion leads to a 60% reduction of NusG engaged with the elongation complex, which we attribute to loss of the anti-termination complex on rRNA operons, subsequent termination, and loss of NusG as elongation complexes dissociate. This interpretation is further supported by our super-resolution analysis (Fig. 3C), which showed that *nusB* deletion leads to a general decompaction of the nucleoid and a loss of the large NusG clusters located in the nucleoid periphery, where the *rrn* operons are expected to reside (Cabrera and Jin, 2006; Gaal et al., 2016). We have also visualized these *rrn* anti-termination NusG clusters directly by halting translation initiation with kasugamycin. Our results establish that, at moderate growth rates, most NusG molecules on the chromosome are involved in translation-independent activities, and specifically in rRNA antitermination. This NusG fraction engaged in *rrn* antitermination is expected to increase further in cells grown in richer media, such as LB and RDM.

The reduction of NusG engagement with the elongation complex is even more dramatic (~90%) when both *rrn* anti-termination and transcription-translation coupling are eliminated by removing the NusG-NusE interactions (using the NusE degron). This comparison strongly suggests that the involvement of NusG in transcription-translation coupling *in vivo* is substantial, with coupling occupying the second largest fraction of the NusG pool after *rrn* anti-termination.

### NusG molecules involved in *rrn* anti-termination reside in phase-separated condensates

The *nusB* deletion (Fig. 3B) reduced the bound NusG fraction, and, intriguingly, yielded an even slower VS-NusG population at ~0.1 μm^2^/s. This striking result suggested that the bound population in unperturbed cells was a convolution between a DNA-bound state, and a state that had some limited mobility and exhibited confined diffusion, as was shown for RNAP and NusA in the large clusters forming during exponential growth in rich media (Ladouceur et al., 2020; Stracy et al., 2015). This same work proposed that RNAP clusters involved in *rrn* antitermination are biomolecular condensates formed via LLPS, and that NusA, a protein with disordered segments, may nucleate condensate formation *in vivo*.

Our observations agree with the Ladouceur *et al*. work and extend its findings. We show that HEX treatment was able to dissolve NusG clusters both in Kas-treated cells (enriched for rRNA transcription and devoid of mRNA transcription), as well as in untreated cells (Fig. 5A). Our results show directly that NusG is part of *rrn* anti-termination condensates in *E. coli*, which we also expect to include NusA and other anti-termination factors. Further, our tracking analysis showed that HEX treatment yielded an almost identical diffusive profile to the one obtained with *nusB* deletion in terms of loss of VS-NusG species and correlated increase of F-NusG species (compare Figs. 5B and 3A), strongly suggesting that *rrn* synthesis occurs almost exclusively in LLPS condensates, providing further strong links between condensate formation and *rrn* antitermination.

### Model of NusG functional allocation *in vivo*

Our results provide a working model for how NusG distributes between its functions in cells to modulate transcription in *E. coli* (Fig. 6). NusG in the cell associates with larger structures and machinery, and is a limiting for NusG-dependent reactions. This landscape creates opportunities for functional modulation of different genes by regulation of NusG concentrations and its complexes in different physiological states.

**Figure 6.**
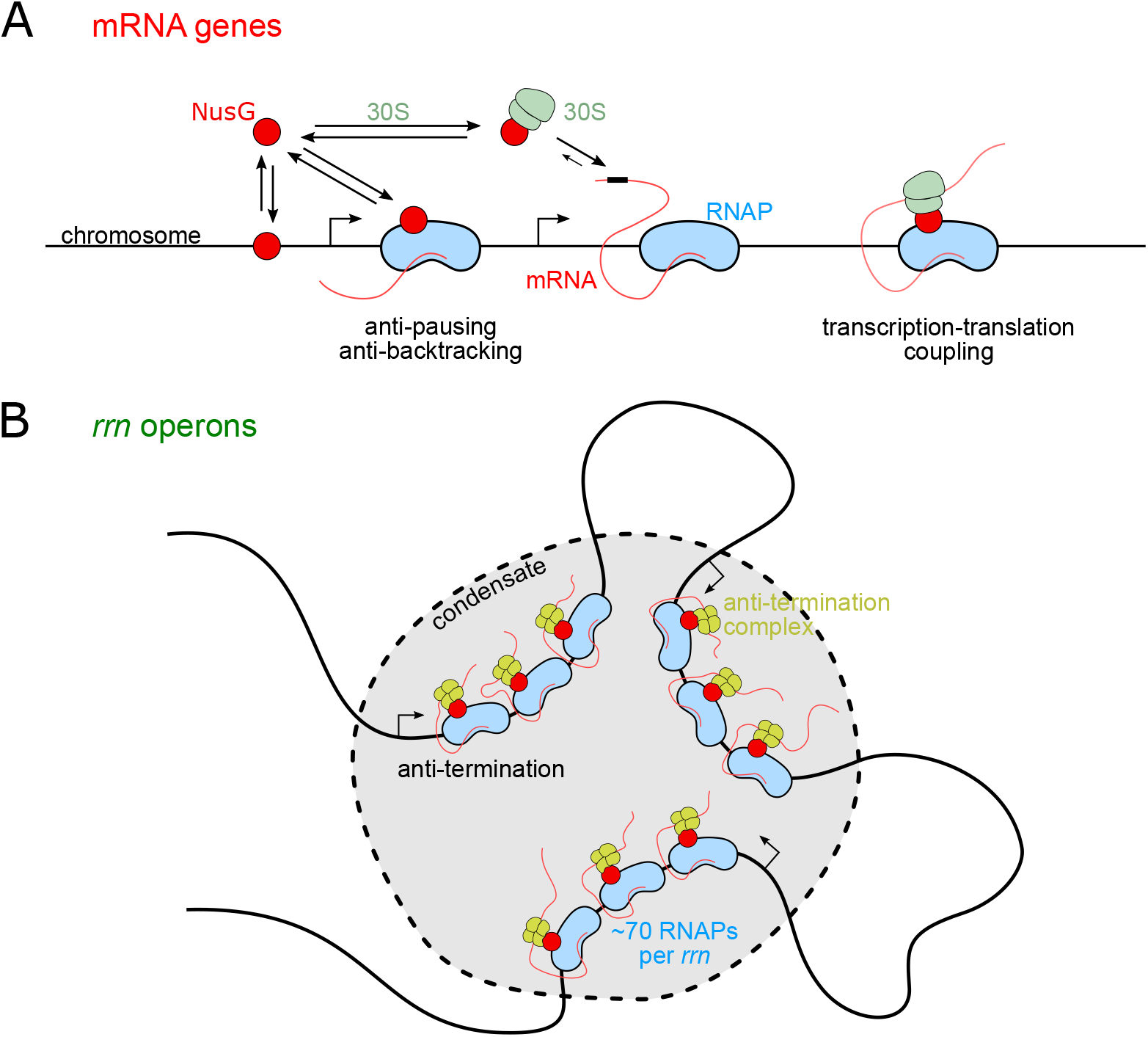
Model of NusG functional distribution inside bacterial cells. **A.** Functional distribution of NusG on mRNA genes. Before locating its target (a transcription elongation complex), NusG interacts non-specifically and transiently with the bacterial chromosome. NusG can enter the elongation complex by binding to RNAP directly. NusG also forms an abundant complex with free 30S ribosomal subunit, which can interact with the mRNA during translation initiation and offer a route for locating the elongating (or paused) RNAP, and establishing transcription-translation coupling. **B.** Role of NusG in rRNA transcription. *rrn* operons are heavily transcribed in moderate to fast growth rates, with each *rrn* operon being occupied by tens of RNAP molecules (~70 for exponential growth in rich media). NusG forms part of the *rrn* anti-termination complex that ensures fast and complete synthesis of rRNA. The *rrn* operons are in close proximity in 3D-space and form part of anti-termination transcriptional condensates forming via LLPS. Most NusG molecules during moderate-to-fast growth rates are occupied in these large transcriptional assemblies.

On mRNA genes (Fig. 6A), NusG can enter the elongation complex by binding to RNAP directly. NusG also forms an abundant complex with free ribosomal subunit 30S, which can interact with the mRNA during translation initiation and offer a route for locating elongating or paused RNAPs, and establishing transcription-translation coupling. Disruption of the NusG-ribosome interaction affects NusG-dependent coupling and leads to loss of NusG from the transcription elongation complex, which may lead to transcription termination.

On rRNA genes (*rrn* operons; Fig. 6B), which are heavily transcribed during moderate to fast growth, NusG forms part of the *rrn* anti-termination complex that ensures that all RNAPs achieve rapid and complete synthesis of rRNA. The *rrn* operons are in close proximity in 3D-space and form part of anti-termination transcriptional condensates generated via LLPS. Most NusG molecules under moderate to fast growth rates are occupied in these large transcriptional assemblies. Our data also suggest that a significant fraction of NusG molecules in the condensates displays confined diffusion within the condensates (as also shown for RNAP), and may be recycled within the condensates upon completion of rRNA synthesis. Regulation of the condensate stability will affect NusG functions, potentially offering powerful means to regulate rRNA transcript levels.

## ACKNOWLEDGMENTS

The authors thank Dr. Piers Turner for providing help with cell segmentation; Dr. Olivier Espeli for providing plasmids; Prof David Sherratt for providing reagents for the NusE degron strain; Prof Robert Landick for providing a helper plasmid; and the MICRON Advanced Bioimaging Facility at Oxford (supported by Wellcome Strategic Awards 091911/B/10/Z and 107457/Z/15/Z) for access to a PALM and a 3D-SIM microscope in their facilities.

## AUTHOR CONTRIBUTIONS

A.N.K. and M.G. conceived the project. A.N.K., M.G, M.S., and H.e.S. designed experiments. H.e.S., M.S., and O.J.P. performed experiments and analysed data. O.J.P. performed simulations, and provided software. A.N.K. analysed data. H.e.S., O.J.P., M.G., and A.N.K. discussed and interpreted data. H.e.S., O.J.P. and A.N.K. prepared figures. H.e.S. and A.N.K. wrote the paper, and all authors had the chance to read and edit the paper.

## DATA AVAILABILITY

Movies and images of cells as well as localisation files for single molecules are available upon request.

## MATERIALS AND METHODS

### Strain construction

NusG-sfGFP and PAmCherry fusions were constructed as N-terminal fusions using gene doctoring (Herring et al., 2003; Lee et al., 2009) as previously described by (Mooney et al., 2009b). Briefly, we used Gibson assembly (Gibson et al., 2009) to clone either sfGFP or PAmCherry coding sequences followed by a flexible linker sequence (GGSGGGSGA) between the start ATG codon and the second codon, flanked by 1-kb homology to serve as the recombination donor.

For the NusE-mNeonGreen-degron tag construction, we had to address the fact NusE is essential and its encoding gene *rpsJ* is the first of 11 genes in a ribosomal protein operon. The DAS+4 degron tag (McGinness et al., 2006) was introduced using the lambda red system (Datsenko and Wanner, 2000) at the end of *rpsJ*, while introducing only the *kan^R^* sequence and an RBS upstream in a way that the *kan^R^* gene behaves as part of the ribosomal protein operon (rather than a standalone insulated cassette).

All strains used in our study are found in Table S1.

### Immunoblotting

Cultures of a strain carrying NusG-PAmCherry and a strain carrying free PAmCherry under the control of a pBAD promoter were diluted from an overnight culture in LB until the OD_600_ reached 0.2; subsequently, 5 ml were collected and spun down, and the pellet was re-suspended in 100 μl of 1xLaemmli buffer. For the arabinose-inducible strain, 0.2% final arabinose was added to the culture, and was left for 20 min before the cells were pelleted and harvested. The cells were boiled for 10 min at 95°C, and 10 μl were loaded on a pre-cast 4-20% mini Protean gel (Biorad) next to 5 μl of pre-stained protein standard ladder. Once migration is completed, western blotting was performed using the P3 program of the iBlot system (Thermo Fisher Scientific). The membrane was blocked using TBST+6% skimmed milk, incubated with Anti-mCherry antibody (ab183628-100μl) in 1:2,000 TBST-milk, and left shaking at 4°C overnight. We then incubated at a ratio 1:5,000 with the secondary antibody of goat anti-rabbit coupled with HRP (A6154-1ML) for 1 hr at room temperature. Finally, the Pierce ECL plus western reagent was added to the membranes according to supplier’s recommendation and imaged used a Typhoon FLA 9500 gel scanner (GE Amersham).

### Growth rate measurements

The strain carrying NusG-PAmCherry and the MG1655 strain were serially diluted in LB 1:1,000, incubated at 37°C in a Clariostar plate reader (GMB) and run overnight with measurements taken every 5 min for up to 24 hours. For doubling time calculations, cells were diluted from an overnight culture in fresh media, with samples were taken every 30 min until reaching an OD_600_ higher than 1.

### Cell preparation for imaging

Single colonies from a streaked plate of the strains were inoculated in one of three media as necessary in this work: M9GluVA (M9 media supplemented with MEM amino acids, MEM vitamins and 0.2% glucose), M9Glu (M9 media supplemented 0.2% glucose and EZ rich defined media) and RDM EZ Rich Defined Medium Kit, without Methionine (Teknova M2125). Cells were grown overnight at 37°C. Cultures containing plasmids were supplemented with the suitable antibiotics of 100 μg/ml ampicillin, or 50 μg/ml kanamycin, or 35 μg/ml chloramphenicol. Overnight cultures were diluted and grown for >2 hr at 37°C to exponential phase (OD_600_<0.2). Cells were centrifuged and immobilized on 1% low-fluorescence agarose (1613100, Biorad) pads, sandwiched between two glass coverslips (no. 1.5 thickness; prior to use, coverslips were heated to 500°C in a furnace for 1 h to remove any fluorescent background particles). Measurements were performed at 21°C. For treatments with inducers or antibiotics, cells were incubated with either 50 μg/ml rifampicin, or 100 μg/ml chloramphenicol, or 500 μg/ml kasugamycin, or 1 mM IPTG for 30 min unless otherwise indicated.

### Single-molecule imaging and tracking

All single-molecule tracking PALM and brightfield (BF) images were acquired using a custom-built microscope, equipped with three lasers, a 200 mW 405 nm diode laser (MDL-III-405, CNI, Changchun, China), a 70 mW 473 nm diode laser (Stradus 473, Vortran, Roseville, CA, USA) and a 200 mW diode-pumped solid-state 561 nm laser (561L-COL-PP, Oxxius, Lannion, France). The lasers were modulated using a DAQ system (NI cDAQ-9274 chassis, NI 9263 module; National Instruments, Austin, TX, USA), with the 405-nm and 473-nm lasers modulated directly via analogue voltage commands, and the 561-nm laser using an acousto-optic modulator (Gooch & Housego, Ilminster, Somerset, UK). A custom-built LabVIEW virtual instrument (National Instruments, Austin, TX, USA) software was written for laser modulation. The lasers were coupled into single mode optical fibres, collimated, reflected by a multi-bandpass optical filter (69013m, Chroma Technology Corp, Bellows Falls, VT, USA), and focused by an achromatic doublet lens (AC508-300-A, ThorLabs, Newton, New Jersey, USA) onto a single point on the back focal plane of a 100x NA1.40 oil immersion microscope objective (UPlanSApo, Olympus). Exit angle modulation enabled sample illumination in epifluorescence, variable angle epifluorescence microscopy (VAEM) and total internal reflection (TIR) modes – for our cellular imaging, only VAEM configuration was employed. BF images were illuminated using a white LED light source (CoolLED pE-100). Collected light was passed back through the multinotch filter with transmission windows at 439±15 nm, 521±17 nm, and 605±25 nm, with an achromatic doublet lens (AC508-300-A, ThorLabs, Newton, New Jersey, USA) forming an image on an EMCCD camera (iXon 897 Ultra, Andor Technology Ltd, Belfast, UK). Image acquisition was performed using the software package Andor SOLIS (Andor Technology, Belfast, UK). Tracking PALM comprised 561-nm excitation at ~340 W/cm^2^, and 405-nm excitation at 0-1 W/cm^2^; imaging parameters involved a pre-amplified gain of 1, a memory parameter of 1, and 10-ms exposures over 30,000 frame recording.

### SIM imaging

For 3D imaging, 3D-Structural Illumination Microscopy (3D-SIM) was performed described (Mamou et al., 2022; Stracy et al., 2015) with minor adjustments. Briefly, imaging was performed using a Deltavision OMX-SR microscopy system (GE Healthcare) equipped with four laser lines (405, 488, 568 and 640 nm), pco.edge 4.4 sCMOS cameras (PCO) and a 60x oil-immersion objective (Olympus PlanApo 1.42 NA). An area of 512×512 pixels was used to acquire a stack of 125-nm sections to generate a total of 2-3 μm thickness. Each z section results from a striped illumination pattern rotated to the three angles (−60°, 0°, +60°) and shifted in five phase steps. NusG-sfGFP was excited using 10% of the 488-nm laser power and imaged using 10-ms exposures, whereas the DAPI stain was excited using 20% of the 405-nm laser power and imaged using 20-ms exposures. The image stacks were 3D-reconstructed using Deltavision softWoRx 7.2.0 software with a Wiener filter of 0.003 using wavelength-specific experimentally determined OTF functions. Average intensity and 3D projections of 3D-SIM images were generated using ImageJ to generate the two-colour 3D-SIM images.

### Image and data analysis for PALM imaging

Image and data analysis were performed as we previously described (Stracy et al., 2015). Briefly, single-molecule localisations in cells were performed using custom in-house tracking software (“StormTracker”). Initial cell segmentation using a segmentation algorithm (based on Mask R-CNN (He et al., 2020)and training on bright-field images) was followed by mesh refinement via Microbetracker (Sliusarenko et al., 2011). Custom in-house software (“LoColi”) was used to filter localisations using a segmentation mask and generate tracks that were used to calculate diffusion coefficients D* for each track. D* histograms were generated from the compilation of triplicate datasets and were fitted to gamma distributions (Stracy et al., 2015). Heatmaps for the intracellular locations of each localisation in the selected tracks were computed for both different diffusive populations (using a specified D* threshold to examine tracks of different mobility), and normalised across cell length.

### Estimation of the fraction of time NusG spends in transient DNA binding

The fraction of time that a DNA-binding protein binds non-specifically to the chromosome can be estimated by a simplified version of (Elf et al., 2007) using D_intact_ = (1-f_ns_) * D_free_, where D_intact_ and D_free_ are the accurate protein diffusion coefficients in cells with intact nucleoids, and in cells in the absence of DNA binding, respectively, and f_ns_ is the fraction of time that the protein binds non-specifically to the DNA.

