## Supplemental Figures for "Single-molecule tracking reveals the functional allocation, *in vivo* interactions and spatial organization of universal transcription factor NusG"

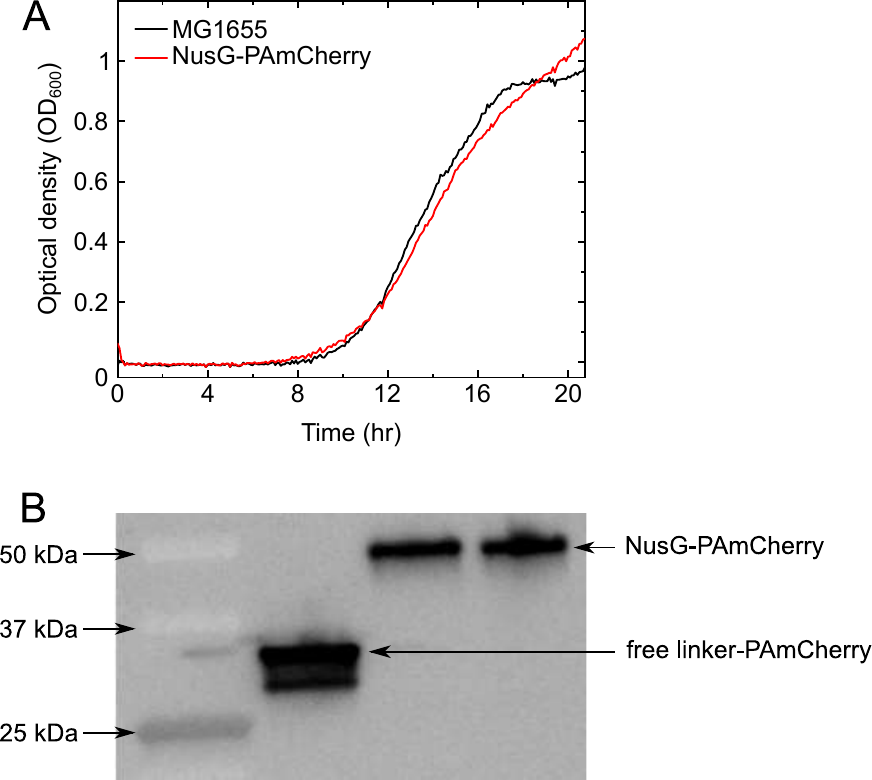

**Figure S1**. **Verification of the NusG-PAmCherry fusion**.

**A.** Growth curve for wild type MG1655 (blue) and the NusG-PAmCherry fusion (red) grown in LB overnight, then diluted 1:1,000 in a 96-well plate in triplicates. Readings were taken every 4 min for 24 hrs.

**B.** Western blot to validate that the NusG-PAmCherry fusion is expressed intact. Left, molecular weight marker; middle, lysate from a control strain DH5α carrying an arabinose-inducible plasmid expressing PAmCherry plus a linker; right, lysate from the genome-encoded NusG-PAmCherry fusion strain. Lysates were probed using the anti-mCherry antibody as described in *Methods*. The intact nature of NusG-PAmCherry is indicated by the band appearing at the combined size of both proteins in comparison to the free PAmCherry fusion alone.

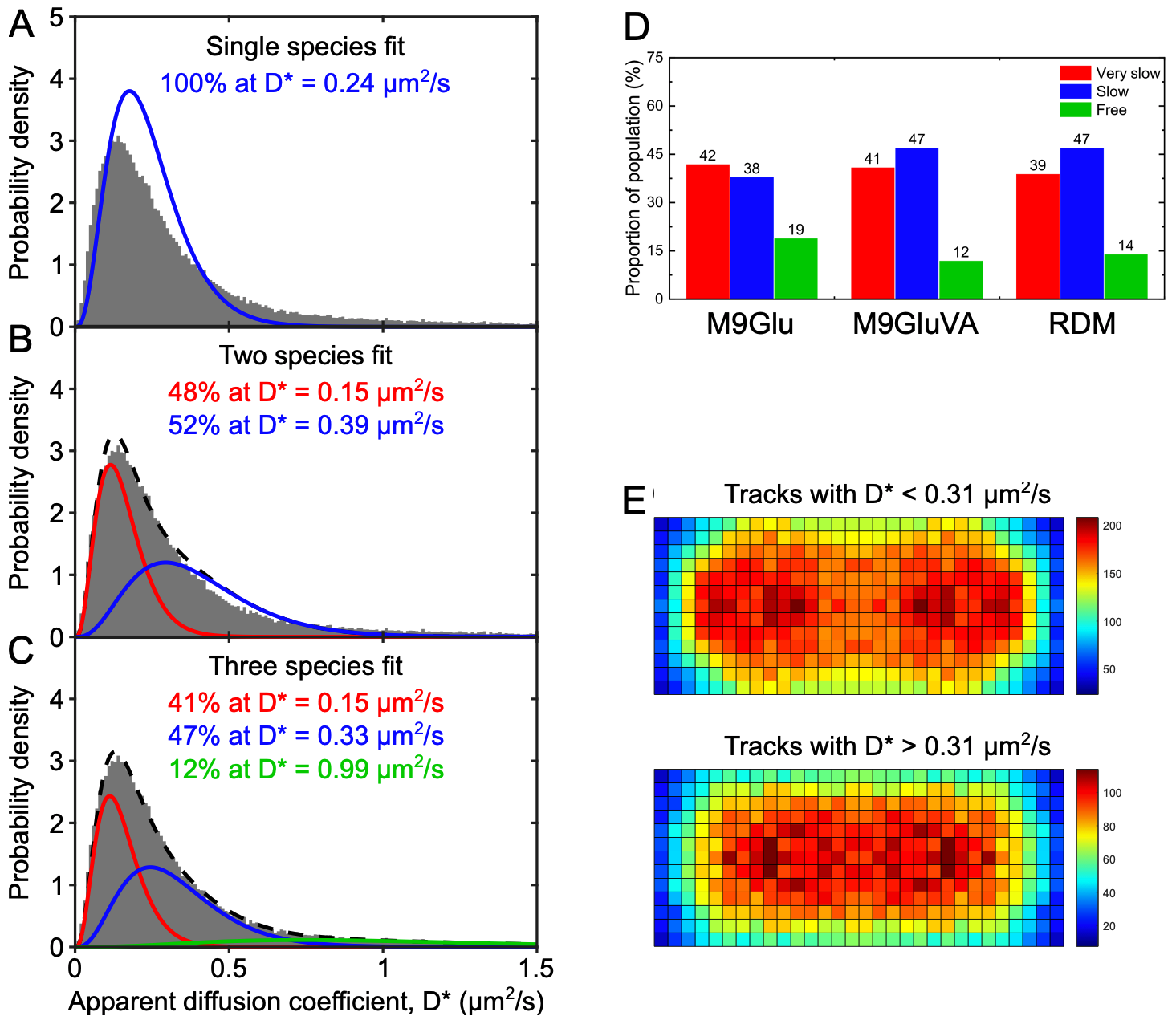

**Figure S2**. **Sorting of NusG D* distribution into different diffusive species, and visualizing their spatial distribution.**

**A.** Single-species fit to the D* distribution of NusG.

**B.** Two-species unconstrained fit performs better.

**C.** Three-species fit performed as described in the main text.

**D.** The fractions of the NusG D* distribution that correspond to the VS-NusG, S-NusG, and F-NusG species for the three growth media used in this study.

**E.** Spatial distribution heatmaps of NusG tracks with a categorization threshold of D* = 0.31 for 244 cells with lengths of ~1-2 μm, having ~1 nucleoid. Top: a heatmap for molecules with D*<0.31 μm²/s, representing mainly the VS-NusG species. Bottom: a heatmap for molecules with D*>0.31 μm²/s, representing mainly the S-NusG and F-NusG species.

**Table S1.** List of bacterial strains and plasmids used in this study.

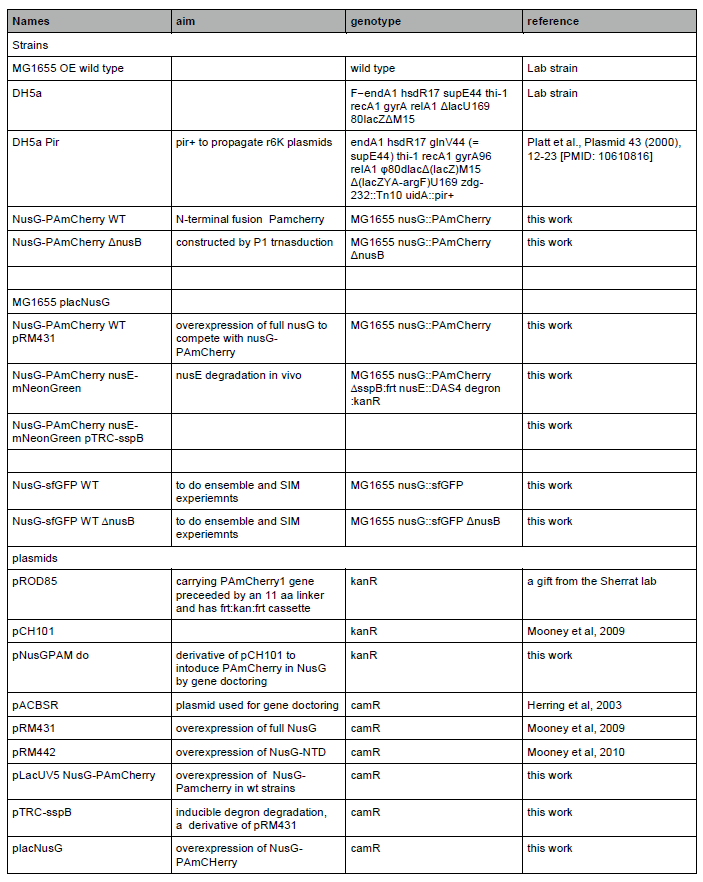
